# The Effect of Neurodegeneration on Ultrasonic Vocalisations (USV) and Their Neuronal Substrates in Mice and Rats: A Systematic Review

**DOI:** 10.64898/2026.02.27.708197

**Authors:** Charissa Calemi, Rose Bruffaerts, Tommas Ellender

## Abstract

This systematic review examines the effects of neurodegeneration in rats and mice on ultrasonic vocalisation (USV) production and its underlying neuronal substrates. Neurodegenerative diseases, such as Parkinson’s, Alzheimer’s, and frontotemporal degeneration, significantly impair communication abilities in humans. Animal models, particularly rats and mice, are widely used to study the underlying mechanisms of these disorders. One important aspect of neurodegeneration is its impact on ultrasonic vocalisations in rodents. USVs play a crucial role in social interaction, mating, and distress signalling, making them valuable behavioural biomarkers for neurological dysfunction. This review aims to synthesise existing research on how neurodegeneration affects USV production and its neuronal substrates in rodent models. Understanding these changes can provide insights into disease progression and facilitate the development of early diagnostic tools and therapeutic strategies. Studying USV impairments in animal models may help identify biomarkers relevant to human speech deficits in neurodegenerative diseases. By bridging the gap between preclinical and clinical research, this review contributes to the growing field of neurobehavioral biomarkers, which could ultimately improve early diagnosis and intervention in human neurodegenerative conditions.

## Introduction

Impairments in speech represent a prominent and often early observed feature of several neurodegenerative disorders, including Parkinson’s disease (PD), Huntington’s disease (HD), Alzheimer’s Disease (AD), amyotrophic lateral sclerosis (ALS), and frontotemporal dementia (FTD). Given the urgent need for cost-effective, scalable methods to detect neurodegenerative diseases at early stages, considerable work has focused on speech as a non-invasive early marker. Despite promising findings, however, a robust understanding of the neurobiological mechanisms underlying the onset and progression of vocal dysfunction remains limited. Changes in speech production are ubiquitous in patients with neurodegenerative diseases. In PD, a 𝛼-Synucleinopathy (Bloem et al., 2021), up to 89% of individuals develop hypokinetic dysarthria, a motor speech disorder that substantially impairs communication (Ho et al., 1998; Logemann et al., 1978; Pinto et al., 2004). In HD, a polyglutamine-expansion disorder caused by mutant huntingtin protein aggregation (Walker, 2013), speech alterations often occur early in the disease course and may precede other clinical manifestations during the prodromal stage (Chan et al., 2019). In AD, an amyloid-𝛽 and tau proteinopathy (Scheltens et al., 2021), linguistic changes are often seen as core disease-related changes reflecting a semantic memory deficit, but changes in temporal and acoustic speech parameters have also been reported (Meilan et al., 2014). FTD encompasses a heterogeneous spectrum of clinical phenotypes, presenting with variable degrees of speech and language impairment (Moore et al., 2020; Rohrer et al., 2015), often characterised by changes in fundamental frequency (Nevler et al., 2017). Within the FTD spectrum, most patients exhibit underlying tauopathy or TDP43-proteinopathy (Mackenzie & Neumann, 2016). Motor speech impairments are hallmarks of tauopathy, spanning phenotypes like progressive supranuclear palsy (PSP) and corticobasal syndrome (CBS). Characteristic changes in tauopathies include apraxia of speech and, not infrequently, dysarthria (Peterson et al., 2021). In ALS-FTD, a TDP-43 proteinopathy (Neumann et al., 2006), a myriad of speech and language abnormalities have been reported (Rusina et al., 2021). Changes in fundamental frequency and different types of dysarthria frequently occur in patients with ALS–FTD (Nevler et al., 2020).

Only relatively recently has the field of neurodegenerative diseases moved to a more quantitative approach to speech changes as part of digital markers. This shift facilitates not only early diagnosis but also longitudinal monitoring in the context of clinical trials (Berthier et al., 2026). Interestingly, a recent systematic review of the FTD spectrum demonstrated that acoustic features such as speech rate, articulation rate, and pause frequency are generalisable across languages, enabling cross-linguistic comparisons (Coppieters et al., 2024). This prompts the question of whether some core changes in neurodegenerative disease are so generic that they may as well apply to other species. In this regard, studies in songbirds such as the zebra finch have been highly informative as a model for human speech (Konopka & Roberts, 2016; Mooney, 2020; Zhang et al., 2023). However, despite songbirds providing an ethologically relevant model of vocal learning, mice offer unparalleled genetic accessibility for causal interrogation of behavioural and pathophysiological aspects of neurodegeneration. They allow for experimental control, the study of neurobiological changes in the preclinical and prodromal stages, and the integration of behavioural outcomes with neurochemical and neuropathological findings, and, in some studies, have also analysed their vocalisations.

Rodents, including mice and rats, emit ultrasonic vocalisations (USVs) in the frequency range above 20 kHz (Anderson, 1954; Wohr & Schwarting, 2013). Although best characterised in rats, USVs can be classified into three broad categories: aversive calls, positive affective calls, and pup calls elicited by maternal separation (Portfors, 2007). Adult rats emit low-frequency (∼22 kHz) USVs in response to aversive stimuli (Hamed et al., 2009; Tornatzky & Miczek, 1994) and high-frequency (∼50 kHz) USVs during rewarding contexts such as mating or play (Panksepp & Burgdorf, 2003). In adult mice, low-frequency calls (≤60 kHz) occur during free movement and restraint, whereas higher-frequency calls (>60 kHz) are typically produced during social interactions (Lefebvre et al., 2020). Due to developmental differences in the pup larynx, rodent pups emit higher-frequency distress calls (∼40-60 kHz) during maternal separation (Bolukbas et al., 2020; Shair et al., 2015; Yin et al., 2016). Multiple acoustic features can be extracted from USV spectrograms, including call number, duration, intensity (dB), bandwidth or frequency range (Hz), peak frequency, pause duration, call rate (calls per time period), and latency to first call. Furthermore, several classification systems have been proposed to categorise USVs into discrete call types based on complexity and frequency modulation (Ahrens et al., 2009; Ciucci et al., 2009; Ey et al., 2013; Holy & Guo, 2005; Johnson et al., 2013; Portfors, 2007; Scattoni et al., 2008; Wright et al., 2010). The two main categories of USVs are simple and complex calls. Simple calls are flat, with a near-constant frequency, while complex calls contain two or more directional changes in frequency; however, no universally accepted framework currently exists.

It is thought that USV production relies on the interplay between the forebrain, midbrain, brainstem, and laryngeal structures. A key region is the midbrain periaqueductal gray (PAG), which acts as a vocal gating centre (Jurgens, 1994, 2002, 2009). Lesions of the PAG result in mutism (Esposito et al., 1999; Jurgens, 1994, 2002, 2009), whereas optogenetic activation of glutamatergic USV-related PAG neurons elicits vocalisations in the absence of social cues (Tschida et al., 2019). The PAG projects to motor and respiratory nuclei in the brainstem, including the nucleus tractus solitarius (NTS), whose disruption leads to complete mutism in genetically modified mice (Hernandez-Miranda et al., 2017). The NTS contains premotor neurons projecting to spinal (L1) motor neurons and the nucleus retroambiguus (nAmb), which control expiratory and laryngeal musculature (Hernandez-Miranda et al., 2017). Additional midbrain and forebrain structures modulate vocal behaviour. The ventral tegmental area (VTA), a key component of the mesolimbic dopamine system, plays an important role in reward-related vocalisation, with lesions significantly reducing frequency-modulated (FM) 50-kHz USVs in rats (Burgdorf et al., 2007). Furthermore, activation of vesicular GABA transporter (Vgat)-expressing neurons in the medial preoptic area (mPOA) promotes USV production in the absence of social stimuli (Gao et al., 2019), whereas neurons at the interface of the central and medial amygdala suppress vocal output (Michael et al., 2020). Cortical regions, including the anterior cingulate cortex and posterior prelimbic cortex, also contribute to vocal initiation and modulation, with neuronal activity in these areas preceding vocal onset (Bennett et al., 2019; Gan-Or & London, 2023). Finally, it was also shown that the primary and secondary motor cortex (M1 and M2), together with the subjacent anterodorsal striatum, were active during vocalisation (Arriaga et al., 2012).

In this systematic review, we explore the literature on USVs in rodent models of neurodegeneration. First, we report on the different experimental designs used to elicit USVs and then compare USV changes across rodent models of neurodegeneration in relation to observed changes in the underlying neurobiology.

## Methods

This review was prospectively registered in PROSPERO (ID: CRD420250615249) and followed the Preferred Reporting Items for Systematic reviews and Meta-Analyses (PRISMA) guidelines (Page et al., 2021). The detailed search strategies are provided in Supplementary Material **S1**. Searches were run on 16 January 2025 using online databases PubMed, Web of Science, and Embase. No date or language filters were applied to the results.

### Screening

Our search yielded 4690 studies. The results were downloaded to citation management software (EndNote version 21) and underwent semi-automatic de-duplication. Next, records were uploaded to the screening platform Rayyan (https://www.rayyan.ai/) for additional deduplication and to screen articles. Following deduplication, 3649 titles and abstracts were screened for eligibility using the prespecified inclusion and exclusion criteria stated below. After abstract screening, 47 reports were assessed for eligibility based on full-text screening. Upon further reading and data extraction, 33 papers were selected for inclusion in the final paper. The full screening process is presented in the PRISMA flowchart (**Figure 1**).

**Figure 1:**
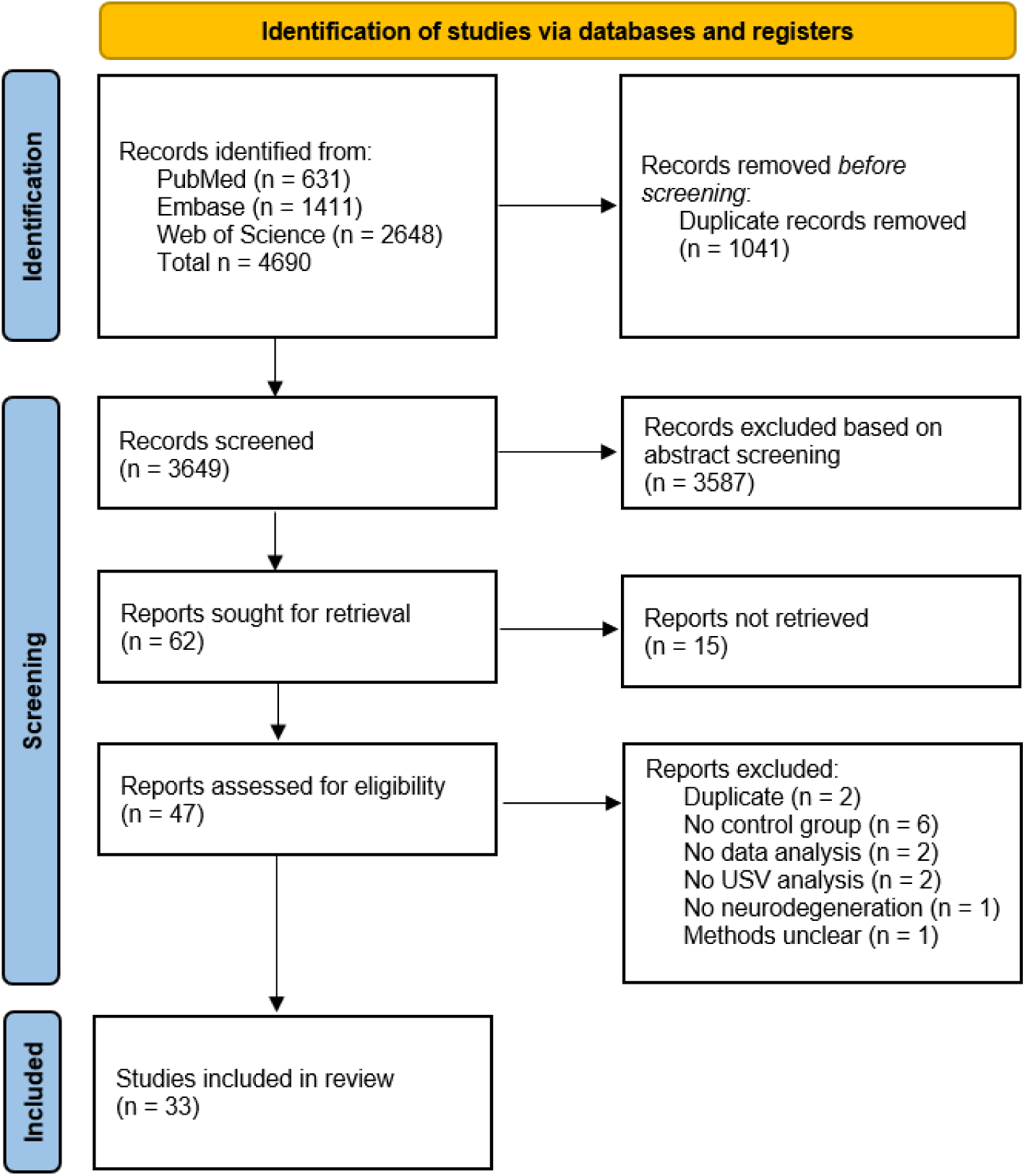
PRISMA 2020 flow diagram for systematic reviews (Page et al., 2021).

### Inclusion criteria

- **Animals/population:** laboratory rats and mice

- **Intervention(s) or exposure(s):** interventions used to induce a model of neurodegenerative disease, such as a genetic model, neurotoxin model, overexpression models, or drug-induced models.

- **Comparator(s) or control(s):** age-/gender-/species-matched wild-type animals

### Exclusion criteria

- **Animals/population:** other laboratory animal models, *ex vivo*, *in vitro*, or *in silico* models, humans

- **Comparator(s) or control(s):** any control receiving a therapeutic intervention

- **Other selection criteria or limitations applied:** (systematic) review articles, meta-analyses, abstracts, conference proceedings, unpublished studies (pre-prints)

### Data extraction and synthesis of the results

The following data were extracted from the eligible studies’ texts, tables, or figures.

1. **Animal model:** species, strain, genotype, age, sex, control group, sample size
2. **USV elicitation paradigm:** type, duration, light/dark cycle
3. USV-related outcomes
4. **Neural findings:** analysis, age, related outcomes

## Results

### Study characteristics

Of the 33 reports included in this review, 21 manuscripts discussed rat models, and 14 discussed mouse models (**Figure 2**). In rat studies, USVs were most frequently studied in models of Parkinson’s disease (PD; 17 papers), followed by Alzheimer’s disease (AD; 3 papers) and Huntington’s disease (HD; 1 paper). In the manuscripts using mice as a model organism, USVs were most frequently studied in models of HD (7 papers), followed by tauopathy (3 papers), PD (2 papers), AD (1 paper), and FTD-ALS (1 paper). For HD, the two used models were R6/1 (5 papers) and BACHD (2 papers).

**Figure 2:**
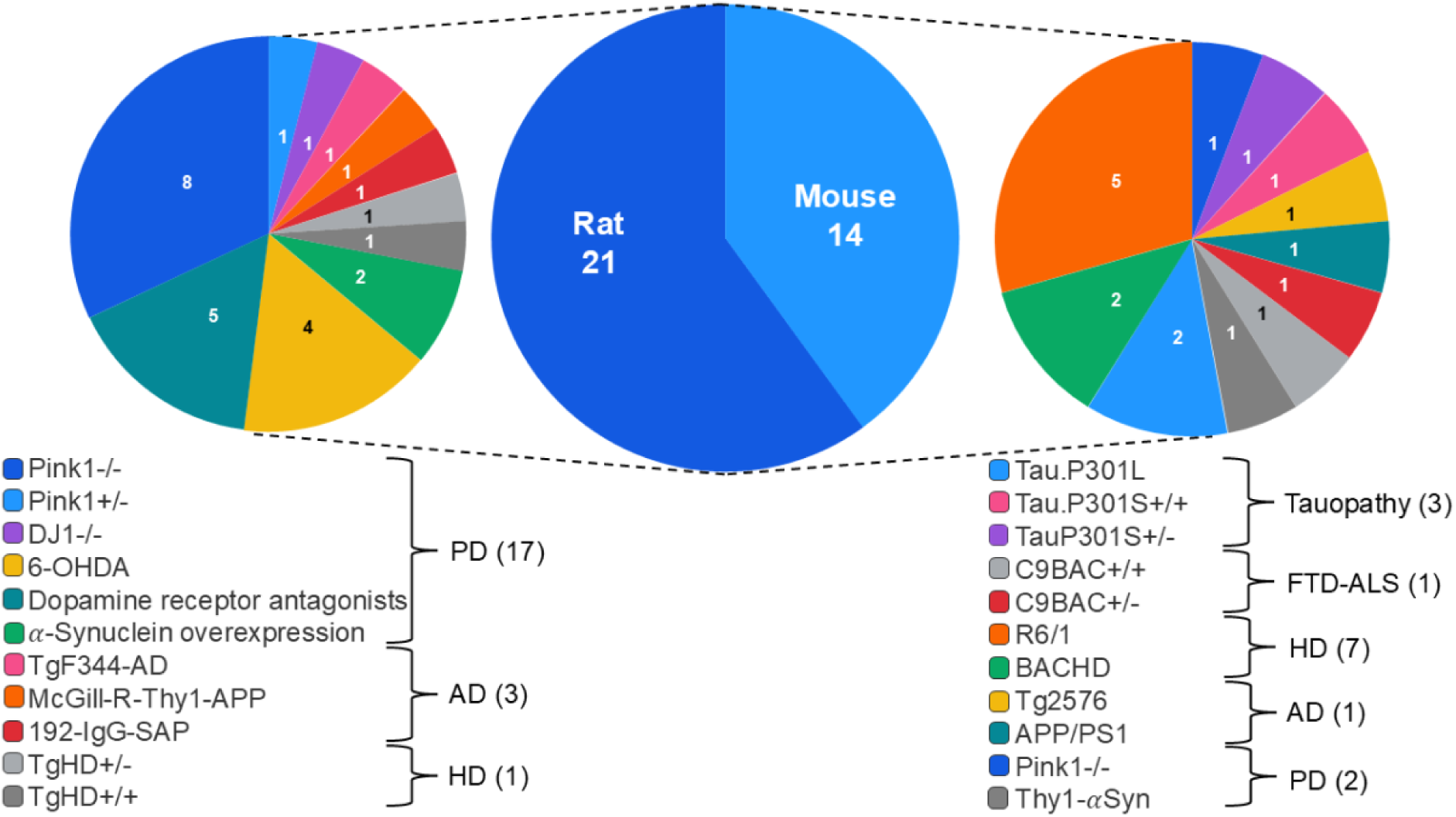
Rodent studies by disease model. Number of peer-reviewed articles published and split by the disease model for rats and mice.

In general, most studies used only males for both species. Of the 21 rat studies, one examined female rats, and one examined both male and female rats. Of the 14 mouse studies, one examined female rats, and five included both sexes.

### USV elicitation paradigms

To record USVs and enable comparisons between neurodegenerative and wild-type animals, experimental paradigms must reliably evoke USVs and often involve either social interaction with a conspecific or the induction of an arousal state in the test animal. Only a limited number of paradigms have been employed across studies (see **Figure 3** for an overview), with courtship being the most frequently used paradigm in both species. In rats (16 papers), the courtship paradigm consisted of the following protocol: a male rat is placed alone in a home cage with a sexually receptive female (in oestrous). After two mounting attempts by the male or after the male demonstrated interest in the female (i.e., sniffing, mounting and/or chasing), the female is removed, and USVs are recorded from the male only (Broadfoot et al., 2024; Converse et al., 2024; Grant et al., 2015; Hoffmeister et al., 2024; Johnson et al., 2020; Lechner et al., 2022; Ringel et al., 2013; Rudisch, Krasko, Barnett, et al., 2023; Stevenson et al., 2019; Yang et al., 2018). Variations on this protocol include putting both animals together for several days until reliable sexual interest is achieved, before recording commences (Gombash et al., 2013; Paumier et al., 2015), or leaving the female in a chamber with ventilation above the male, so the pheromones pass through while recording the male USVs (Ciucci et al., 2007; Ciucci et al., 2008; King et al., 2016). Finally, Marquis and colleagues recorded the female USVs only after removing the male (Marquis et al., 2020).

**Figure 3:**
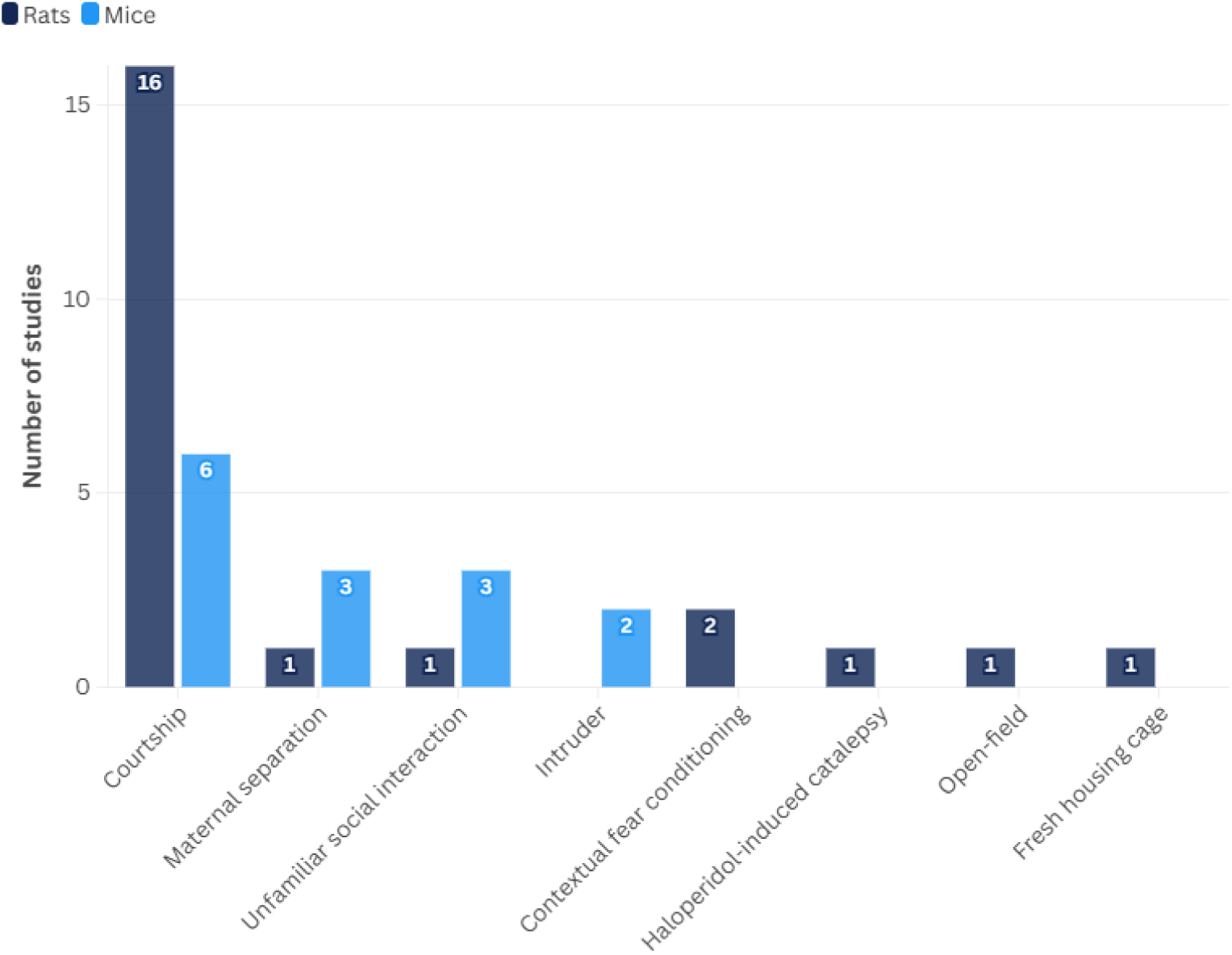
The number of rat and mouse peer-reviewed papers that employ a specific paradigm to elicit USVs.

For mouse studies (6 papers), some papers used the same standard protocols as described for rat studies (Grant et al., 2014; Kelm-Nelson et al., 2018; Mo et al., 2014b). However, there was substantial variability in the courtship paradigm across the mouse studies. For example, two studies used female urine to elicit USVs (Mo et al., 2014a, 2014b, 2015). During this protocol, male mice were first exposed to females in the proestrus or oestrus phase. After female exposure, male mice were exposed to a cotton-tip applicator dipped in urine. The urine collected was from a different female than the one to which it was exposed. Finally, one study did not remove the female, and therefore, USVs were recorded simultaneously from both the male and female mice (Trevizan-Bau et al., 2019).

The second most used paradigms were maternal separation in pups and unfamiliar social interaction in adults. First, in the maternal separation paradigm, both rat (1 paper) and mouse (3 papers) pups were separated from their litter and individually recorded (Cheong et al., 2020; Scattoni et al., 2010; Siebzehnrubl et al., 2018). Secondly, the unfamiliar social interaction paradigm differs across groups and may lean toward the courtship paradigm for some. In rats (1 paper), two male rats from different cages were placed together in an open field (Petrasek et al., 2018). In mice (3 papers), one study placed one male and two female mice of the same age and genotype into the recording cage (Menuet et al., 2011). The animals had no prior interaction and were not sexually naïve. In another study, an unfamiliar, 34-month-old female NMRI stimulus mouse was introduced into the testing cage together with a male mouse (Ragot et al., 2015). The NMRI strain is commonly used in studies of social behaviour because of its high sociability (D’Amato & Pavone, 1996). Finally, the test of Kahriman and colleagues consisted of three phases: habituation, barrier, and interaction (Kahriman et al., 2023). After the habituation period, the experimental mice were exposed to a same-sex, unfamiliar, 12-week-old non-transgenic stimulus mouse, which was introduced behind a wire mesh. Finally, the barrier was removed, and the mice were allowed to interact.

In the intruder paradigm (2 papers), direct social interaction was assessed in the home cage, where the animals had been previously isolated for about 24 hours (Pietropaolo et al., 2011; Pietropaolo et al., 2012). An unfamiliar stimulus mouse (a 12-week-old NMRI female) was then introduced into the testing cage and left there for a few minutes. At the end of the first encounter, the intruder was removed and left in a waiting cage before a second encounter began.

In rats, low-frequency USVs around 22 kHz are part of the rodent defensive repertoire (Brudzynski, 2009). To elicit these specific USVs, the animal is subjected to an aversive stimulus, such as a foot shock, during contextual fear conditioning (FC). The FC encompassed three sessions: training, context and tone. During the training session, a tone is paired with a foot shock. During the context session, rats are placed in the training chamber without any tone or foot shock. In the tone session, rats are exposed to the tone without any shock. This test was performed by Cutuli and colleagues and Colombo and colleagues, although the latter study did not include the presentation of a tone (Colombo et al., 2013; Cutuli et al., 2013).

Finally, USVs were also assessed in additional behavioural contexts, including haloperidol-induced catalepsy and a standard open-field test (Colombo et al., 2013), as well as after the transfer of animals to a fresh housing cage (Vecchia et al., 2018). Neither the open-field test nor the haloperidol-induced catalepsy paradigm elicited USVs. These results show heterogeneity across paradigms used in both species, as well as variability within paradigms. The lack of standardisation complicates comparisons across studies.

### Changes in USV parameters

Hereafter, we describe all significant changes in USV features across the different rat and mouse models. Interestingly, despite the aforementioned variations in USV recording paradigms, we found that similar features underwent significant changes across multiple studies in the same animal model (**Table 1**). Additionally, an overview of all the features analysed across more than one study is summarised in **Table 2**.

**Table 1.**
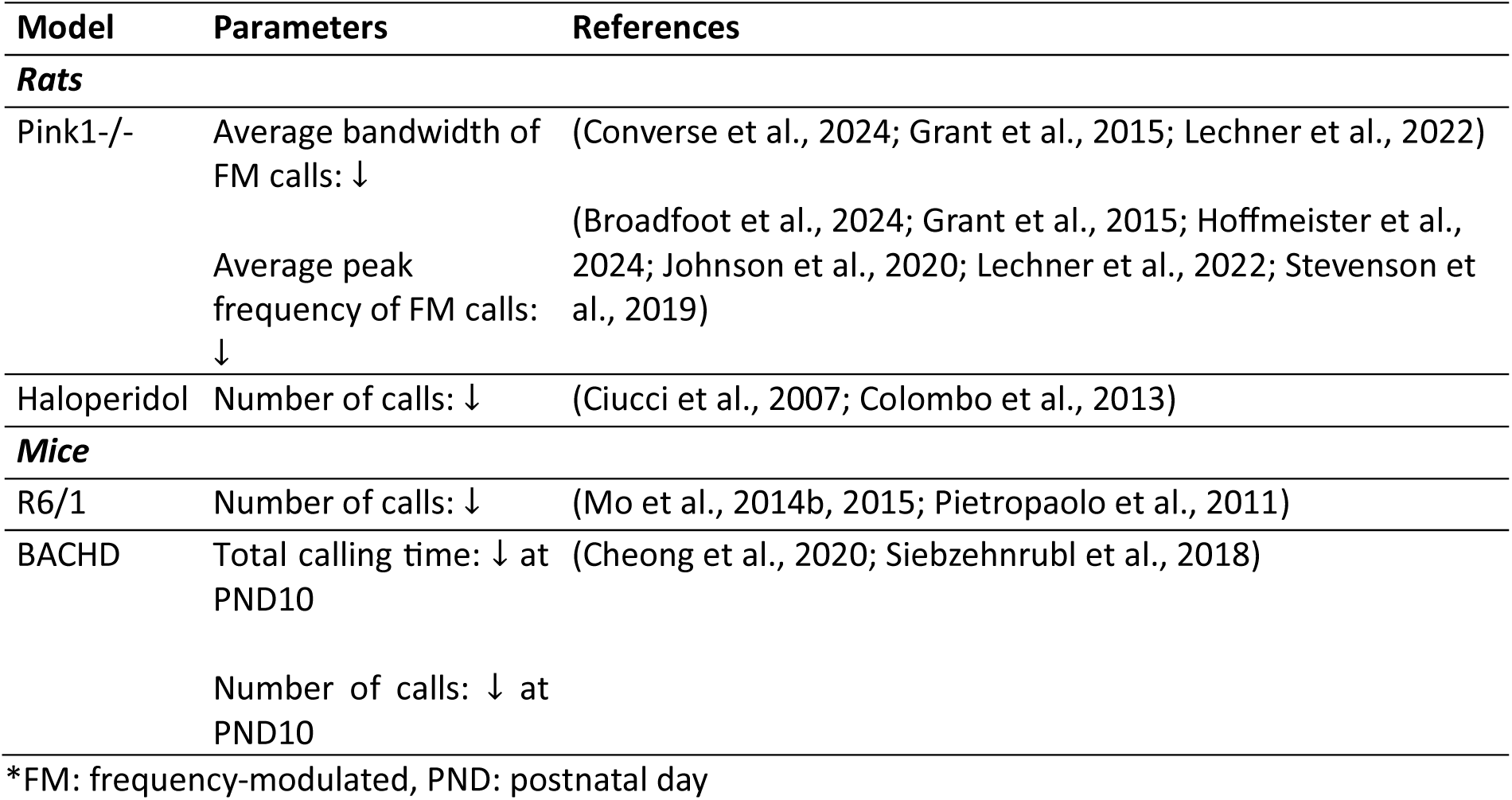
Summary of Significant Vocal Parameters Across Studies Using the Same Animal Model.

**Table 2.** Definitions of Features Studied in More Than One Animal Model.

| **Feature** | **Definition (unit)** |
| --- | --- |
| Call duration | Length of each call (s) |
| Total calling time | Total time spent vocalising within a recording session (s) |
| Latency to call | Time delay before the first call is emitted (s) |
| Call number | Total number of vocalisations produced |
| Call rate | Number of calls per unit time (s) |
| Peak amplitude | Maximum intensity level of a call (dB) |
| Peak frequency | Frequency at the greatest amplitude within a call (Hz) |
| Bandwidth | Difference between highest and lowest frequencies (Hz) |
| Intensity | Intensity level of a call (dB) |
| Intensity range | Difference between maximum and minimum intensity within a call (dB) |
| Mean frequency | Average frequency across a call (Hz) |
| Mean power | Average acoustic power per unit frequency (dB/Hz) |
| Percent complex calls | Percentage of calls classified as complex (%) |
| Amplitude at start/centre/end | Intensity measured at different time points within a call |
| Principal frequency | Median frequency of the frequencies within the call (Hz) |
| Low frequency | Lowest frequency component of a call (Hz) |
| High frequency | Highest frequency component of a call (Hz) |
| Call presence | Proportion of calls emitted within a recording session |

### Parkinson’s Disease models

#### Neurotoxin models

##### 6-Hydroxydopamine

Oxidopamine or 6-hydroxydopamine (6-OHDA) is a neurotoxic drug that selectively and acutely destroys catecholamine-containing neurons (Ungerstedt, 1968). A well-known PD model is produced by the unilateral or bilateral intracerebral injection of 6-OHDA into the nigrostriatal pathway (Berger et al., 1991; Sakai & Gash, 1994; Sauer & Oertel, 1994). Vecchia and colleagues showed that bilateral SNc lesions in the rats significantly reduced the dopamine concentration in the neostriatum compared to the vehicle-treated SHAM group (Vecchia et al., 2018).

In rats, nigrostriatal dopamine depletion induced by unilateral 6-OHDA infusion into the medial forebrain bundle results in significant changes in 50-kHz USVs. For the acoustic parameters, reductions were observed in bandwidth, peak amplitude, and call intensity of trill-like 50-kHz calls (Ciucci et al., 2007; Ciucci et al., 2008). A subsequent study by King and colleagues reported that the duration of low-frequency (20–35 kHz) calls was reduced relative to controls, whereas call bandwidth was increased (King et al., 2016). In addition, both the call rate and the total number of calls were significantly decreased. The authors further observed a marked reduction in vocalisation complexity across all three call types examined: low-frequency calls, high-frequency calls (USVs in the 35–100 kHz range), and complex calls (defined as more than two simultaneously produced USVs).

In agreement with unilateral lesion studies, rats with bilateral substantia nigra pars compacta (SNc) lesions showed a significant reduction in the total number of calls, duration and total time spent producing simple (flat) calls, while latency to initiate calls increased (Vecchia et al., 2018).

##### Dopamine receptor antagonists

Dopamine receptor antagonists bind to dopamine receptors and thereby can mimic the dopamine deficiency seen in PD (Beaulieu & Gainetdinov, 2011). Furthermore, selective antagonists allow blockade of either the D1 or D2 receptor subtypes, thereby enabling investigation of the precise roles of both subtypes in the vocal deficits observed in PD. An often-used D2 antagonist is haloperidol (Sanberg, 1980; Vasconcelos et al., 2003).

Colombo and colleagues showed that the duration and number of 22-kHz USVs were reduced in haloperidol-injected rats during the testing session of the conditioned fear experiment (Colombo et al., 2013). For trill-type 50-kHz USVs at 6 months of age, Ciucci and colleagues reported reduced mean bandwidth (Ciucci et al., 2007) in haloperidol-treated animals compared to controls. Also, the number of USVs was reduced compared to controls and 6-OHDA-treated animals (Ciucci et al., 2007).

In another study, Ringel and colleagues investigated the effects of different dopaminergic antagonists on 50-kHz USVs in rats (Ringel et al., 2013). They reported that there was a significant reduction in call rate, average call duration, and bandwidth, as well as in maximum duration, intensity, and bandwidth, in animals treated with SCH-23390 (D1 antagonist), Eticlopride (D2 antagonist), and their combination (D1+D2), compared to saline controls. These reductions were consistently more pronounced under combined D1+D2 receptor blockade than after selective D1 or D2 antagonism alone. Call latency, on the other hand, was markedly increased after all treatments, with the greatest effect after combined treatment. The proportion of complex calls was reduced only in the combined D1+D2 condition.

Similarly, the average peak frequency was significantly lower with both D1 and D2 antagonists relative to controls, while the maximum peak frequency was reduced following D1, D2, and combined treatments.

Overall, combined D1+D2 receptor blockade produced more severe impairments in USV features than selective D1 or D2 receptor antagonism alone, indicating that both dopaminergic pathways contribute to USV production and modulation.

#### Alpha-synuclein overexpression models

Besides dopaminergic neuronal cell loss, PD pathology is also characterised by the presence of misfolded, fibrillar alpha-synuclein in aggregations (Lewy Bodies) and dystrophic neurites (Spillantini et al., 1998; Spillantini et al., 1997). The involvement of alpha-synuclein in the pathogenesis of PD is clearly implicated from duplication, triplication, or point mutations in the *SNCA* gene in autosomal dominant forms of PD (Ibanez et al., 2009; Polymeropoulos, 1998; Singleton et al., 2003). Three ways to overexpress alpha-synuclein in experimental models are through genetic mutations, viral transmission, or intrastriatal injection of alpha-synuclein.

The first paper looked at the vocal deficits in male rats 8 weeks post-administration of rAAV2/5-human-𝛼-syn injections unilaterally in the SNc (Gombash et al., 2013). The authors found that only call intensity and rate of simple and FM calls were significantly reduced compared to wild-type rats.

Paumier and colleagues studied the effect of unilateral intrastriatal injections of either non-fibrillised recombinant mouse 𝛼-syn or injections of 𝛼-syn pre-formed fibrils (PFF) on the production of USVs in rats (Paumier et al., 2015). Significant changes in USV production were observed 180 days post-injection. Both the maximum value and the mean of the 10 highest call durations were significantly reduced in recombinant 𝛼-syn- and PFF-treated rats as compared to naïve age-matched controls. In addition, both the maximum peak frequency and the mean of the 10 highest peak frequencies were lower in α-syn PFF-treated rats than in controls. Furthermore, the total number of simple calls (constant frequency) and the call rate were reduced in 𝛼-syn PFF-treated rats compared with naïve controls.

One additional study characterised vocal deficits in mice overexpressing human wild-type alpha-synuclein under the Thy-1 promoter (Thy1-𝛼Syn) (Grant et al., 2014). The call profile of Thy1-𝛼Syn mice showed significant differences compared with wild-type animals. The percentage of two-cycle calls was significantly reduced in Thy1-𝛼Syn compared to WT at 2-3 months, while the percentage of jump down calls was increased in Thy1-𝛼Syn compared to WT at 6-7 months. Furthermore, at 2-3 months, the average duration of calls was significantly decreased (for harmonic, jump-down, half-cycle, and cycle calls), and at 6-7 months, the intensity of simple calls was significantly reduced in the Thy1-𝛼Syn group. Call types were visually classified into 9 categories (simple, upsweep, downsweep, jump up, jump down, harmonic, half cycle, cycle, and two cycle), adapted from Portfors’s classification scheme (Portfors, 2007).

#### Genetic models

##### DJ1-/- model

The identification of genetic mutations that lead to inherited forms of Parkinson’s has led to the development of novel animal models of PD (Dave et al., 2014). One of these models is the rodent DJ1-/- model mimicking the loss-of-function mutations in the *PARK7* gene encoding the DJ1 protein, which leads to an autosomal recessive form of inherited PD (Bonifati et al., 2003).

Yang and colleagues are the only published paper that investigated vocal changes in the DJ1-/- rat model (Yang et al., 2018). For each investigated parameter, the authors examined the maximum and average values. The maximum is defined as the animal’s best performance, while the average represents the animal’s performance across the full testing session. For the acoustic parameters, the maximum and average durations of 50-kHz calls were longer in DJ1-/- rats than in wild-type rats. For the average call intensity, a significant reduction in 50-kHz calls was observed in DJ1-/- animals at 8 months of age compared with age-matched wild-type controls. Finally, the percentage of complex calls was significantly higher in DJ1-/- animals than in controls.

##### Pink1-/- model

Most PD cases are sporadic, yet 10% have a genetic cause (Lesage & Brice, 2009). Of these genetic cases, mutations in the *Pink1* gene are the second most common cause of autosomal recessive PD (Krohn et al., 2020). PTEN-induced kinase 1 (PINK1) is a mitochondrial serine/threonine kinase that plays a central role in mitochondrial quality control (Narendra & Youle, 2024). Loss-of-function mutations in PINK1 are a well-established cause of autosomal-recessive early-onset PD (Siuda et al., 2014). The Pink1-/- rat model, in which the Pink1 gene is genetically deleted, has become a valuable experimental model for studying PD-related behavioural deficits (Gispert et al., 2009).

For the PD-like models, the Pink1-/- rat has been most extensively studied with respect to vocal deficits. Within the last decade, eight papers in total discussed the effect of the Pink1-/- mutation on the production of USVs in rats.

Among all published studies using Pink1-/- rat models, two acoustic parameters exhibited similar trends in two or more independent studies. First, three studies described a significant reduction in the average bandwidth of 50-kHz FM calls, defined as USVs exhibiting dynamic frequency changes across time, in 2-month-old (Lechner et al., 2022), 4 and 6 months old (Grant et al., 2015), and between 9 and 11 months old Pink1-/- rats compared to control animals (Converse et al., 2024). A second feature that was affected in five papers was the average peak frequency of FM calls (kHz). The peak frequency was significantly lower at 2 months (Broadfoot et al., 2024; Lechner et al., 2022), 4 months (Broadfoot et al., 2024), 6 months (Hoffmeister et al., 2024), 8 months (Stevenson et al., 2019), and 10 months of age (Hoffmeister et al., 2024; Johnson et al., 2020) in Pink1-/- rats compared to wild-types. These findings provide evidence that the bandwidth and peak frequency of USVs are altered early in Pink1-/- rats and decline over time. Grant and colleagues also investigated the Pink1+/- rat and showed that the average peak frequency of FM calls is reduced at 6 and 8 months in Pink1-/- rats compared to their heterozygous counterparts (Grant et al., 2015). At 6 and 8 months, Pink1+/- rats showed a significant increase compared with wild-type rats.

In addition to these robust and significant changes in bandwidth and peak frequency, other USV features were also affected in Pink1-/- rat models, but less frequently observed.

Marquis and colleagues found that the number of calls was increased in female Pink1-/- rats compared to controls at 2 months of age (Marquis et al., 2020). Interestingly, both genotypes produced more calls during oestrus than during non-oestrus. In addition, Converse and colleagues showed that the mean power of simple and FM calls was significantly reduced in 9 and 11-month-old Pink1-/- rats compared to control animals (Converse et al., 2024).

Finally, for several parameters, contradictory results were found, specifically for intensity (“loudness”). Male (Grant et al., 2015) and female (Marquis et al., 2020) Pink1-/- rats produced FM calls with significantly lower intensity (“quieter”) than wild-type and Pink1+/- rats at 2, 4, 6, and 8 months of age. In contrast to Grant and Marquis, Johnson and colleagues and Lechner and colleagues showed that the average intensity of FM calls was actually increased (“louder”) in Pink1-/- rats compared to wild-type controls at 2 months of age (Lechner et al., 2022) and 10 months of age (Johnson et al., 2020). These contrasting findings are interesting and will be discussed further in the Discussion.

Besides differences in average bandwidth, peak frequency and intensity, Lechner and colleagues also found other vocal deficits in 2-month-old Pink1-/- male and female rats that were not reported by other authors (Lechner et al., 2022). The authors showed that the average percentage of complex calls was higher in Pink1-/- rats than in wild-type rats. The top 10 and average durations of FM calls were shorter in Pink1-/- rats than in wild-type rats. It was also shown that the maximum duration of FM calls was shorter in male Pink1-/- rats than in wild-type rats, but this was not observed in female rats. In addition, the maximum and top-10 bandwidth and peak frequency of FM calls were also reduced in Pink1-/- rats.

The number of studies in mice that have correlated changes in vocalisations with explicit changes in the nervous system is much lower. Kelm-Nelson and colleagues characterised early-onset vocal deficits in a Pink1-/- mouse model (Kelm-Nelson et al., 2018). First, regarding the intensity of simple calls, this paper found that Pink1-/- mice produced calls at lower intensity than wild-type animals. Secondly, the percentage of cycle calls was significantly lower in Pink1-/- mice than in control animals. The percentage of simple calls, on the other hand, was increased at 4 months and decreased at 6 months of age in Pink1-/- mice.

In conclusion, studies regarding PD demonstrate that the Pink1-/- model exhibits early-onset alterations in USV features, particularly in average bandwidth and peak frequency of 50-kHz FM calls (see **Table 1**).

### Huntington’s Disease models

#### Genetic models

##### tgHD

The transgenic HD (tgHD) rat model carries a truncated huntingtin cDNA fragment with 51 CAG repeats under the control of the native huntingtin promoter (von Horsten et al., 2003).

Within the field of HD, only one paper looked at the effect of hemizygous and homozygous tgHD mutations on USVs in 10-day-old rat pups. Both the total duration and the number of calls were reduced in hemizygous and homozygous tgHD rats compared to wild-type rats (Siebzehnrubl et al., 2018).

##### BACHD

The bacterial artificial chromosome transgenic mouse model of HD (BACHD) expresses a full-length human mutant huntingtin (*fl-mhtt*) with 97 CAA-CAG mix repeats under the control of the endogenous *htt* regulatory machinery on the BAC (Gray et al., 2008).

The effects of a 97-glutamine repeat expansion on early vocal development were examined in mouse pups aged 0-10 days postnatally. Two independent studies reported that both call number and total call duration were reduced in BACHD pups at PND10 (Cheong et al., 2020; Siebzehnrubl et al., 2018). In addition, Cheong and colleagues observed a reduction in peak amplitude at PND5, as well as dynamic changes in call complexity: the proportion of multi-component calls was increased shortly after birth but decreased by PND10 in BACHD pups compared with wild-type controls (Cheong et al., 2020). To characterise call structure, Cheong and colleagues categorise vocalisations as either simple (constant, FM, and short) or multi-component (frequency steps and composite calls) (Cheong et al., 2020). Composite calls consist of two harmonically independent components emitted simultaneously, whereas frequency steps are characterised by abrupt, vertically discontinuous changes in frequency on spectrograms without temporal interruption (Scattoni et al., 2008).

##### R6/1

The R6/1 mouse line carries a mutant human exon 1 of the *htt* gene containing an expanded CAG repeat sequence of approximately 115 repeats. (Mangiarini et al., 1996). R6/1 mice are characterised by an adult onset.

Several papers showed that the number of USVs produced at 3 months of age (Pietropaolo et al., 2011) and at 3.5 months of age (Mo et al., 2014b, 2015) was significantly reduced in R6/1 mice compared to controls. Mo and colleagues showed that this reduction was significant only during the first minute of the test, compared with the last minute of the 5-minute test (Mo et al., 2015).

Pietropaolo and colleagues showed that the frequency and duration of USVs were significantly reduced in 3-month-old R6/1 mice in males and in both sexes, respectively (Pietropaolo et al., 2011).

These findings show that HD models exhibit early alterations in USV profiles, spanning both developmental and adult stages. Across transgenic rat and mouse models, reductions in call number and duration emerge early in life, suggesting that mutant huntingtin (disrupts the neural circuits underlying vocal communication during critical periods of neurodevelopment. In adult-onset models such as R6/1, continued deficits in vocal output further indicate that these impairments persist and may worsen with disease progression.

### Alzheimer’s Disease models

#### Neurotoxin models

In one study, the authors injected the immunotoxin 192-immunoglobulin G (IgG)-saporin (Sap) into the medial septum and nucleus basalis magnocellularis of rats, causing a selective depletion of basal forebrain cholinergic neurons, which mimics the neuropathological features and cognitive impairments associated with AD (Waite & Chen, 2001; Wrenn & Wiley, 1998). The authors recorded and analysed 22-kHz USVs during context and tone fear conditioning 2.5 weeks after the surgery (Cutuli et al., 2013). No changes were detected in peak amplitude, peak frequency, frequency modulation, or duration, whereas the number of calls was reduced during the training session. Although the lesion did not alter the freezing response, it decreased the concomitant production of 22-kHz USVs in Sap-lesioned rats (Cutuli et al., 2013).

#### Genetic models

Two different rat models are studied for alterations in their vocal abilities.

##### TgF344-AD

The first model is the TgF344-AD transgenic rat, which expresses human APP with the Swedish mutation (APP K670_M671delinsNL-Swedish) and human PSEN1 with the Δ exon 9 mutation, both driven by the mouse prion promoter (Cohen et al., 2013).

The TgF344-AD rat was studied at 6 and 9 months of age (Rudisch, Krasko, Barnett, et al., 2023). Here, the authors found that the duration of FM calls was increased at 9 months in TgF344-AD rats compared with wild-type rats at 6 months of age. For the average mean power of simple and FM calls, a significant reduction was observed in TgF344-AD rats from 6 to 9 months of age. TgF344-AD animals showed a significant increase in the average high-frequency of simple calls from 6 to 9 months, whereas this was not observed in controls. Finally, the average peak frequency of simple and FM calls was increased from 6 to 9 months in TgF344-AD rats.

##### McGill-R-Thy1-APP

The second model is the McGill-R-Thy1-APP rat model, which expresses human APP_751_ with the Swedish double mutation (K670N, M671L) and Indiana mutation (V717F) under the control of the murine Thy1.2 promotor (Leon et al., 2010).

The McGill-R-Thy1-APP rat model was studied at 7 months of age (Petrasek et al., 2018). This study found that the number of simple high-frequency vocalisations (calls without major steps or trills, corresponding to types 1-4 and 13 in the classification by (Wright et al., 2010)) was reduced in McGill rats compared with wild-type, whereas stepped and composite vocalisations (calls containing notable steps in frequency, or a combination with trills, corresponding to types 6-9 and 14 in the Wright et al.) and trills (high-frequency calls with rapid, periodic oscillations over a wide range of frequencies, corresponding to types 10-12 in Wright et al.) did not differ between McGill rats and controls.

##### Tg2576 and APP-PS1

In the literature of mouse studies, only one paper reported abnormalities in ultrasonic communication in mouse models with AD pathology. In this paper, Pietropaolo and colleagues examined APP (Tg2576) and APP-PS1 mouse models at 3 and 6 months of age, i.e., in the early phases of AD pathology. These transgenic lines harbour the Swedish (K670N: M671L) mutation in the amyloid precursor protein (APP), either alone (the Tg2576 or APP mouse) (Hsiao et al., 1996) or in combination with mutations in Presenilin 1 (PS1; the APPPS1 mouse) (Holcomb et al., 1998). The authors found that the number of USVs produced was reduced in APP and APP-PS1 mice compared with wild-type mice at both 3 and 6 months of age (Pietropaolo et al., 2012).

Collectively, these studies demonstrate that AD-related pathology is associated with alterations in USV patterns across both neurotoxin-induced and genetic models. While selective cholinergic lesions primarily reduced call number without affecting acoustic structure, transgenic rat and mouse models exhibited age-dependent and genotype-specific changes in both quantitative and qualitative vocal parameters. These findings suggest that AD-related neurodegeneration and amyloid pathology can disrupt multiple aspects of vocal communication, although the nature and extent of these effects vary across models, species, and stages of disease progression.

### Models with tauopathy

#### Genetic models

To investigate the mechanism by which tau exerts its effects, various transgenic tau mouse models that are based on mutations linked to human tauopathies have been generated (Gotz & Gotz, 2019; Spillantini & Goedert, 2013). Here, two models are described: Tau.P301L and Tau.P301S mouse models. Both models are driven by frontotemporal dementia with Parkinsonism linked to chromosome 17 (FTDP-17) mutations. The P301S mutation causes early-onset FTDP-17, with rapidly progressive and often severe clinical features (Baba et al., 2007; Bugiani et al., 1999). In the field of tauopathy, three papers examined USVs in mouse models.

##### Tau.P301L

The first two described USVs in Tau.P301L mice at 4-5 and 8-10 months of age (Menuet et al., 2011) and 3-4, 5-6, and 8-9 months of age (Trevizan-Bau et al., 2019). Menuet and colleagues showed that the mean frequency of USVs was lower in the Tau.P301L mice than in the wild-type controls in the youngest age group. More drastic results were observed in the older age group, where the duration, number of USVs per minute, total time of USV production, frequency range, and complexity of USVs produced by the Tau.P301L mice were decreased (Menuet et al., 2011).

Trevizan-Bau and colleagues used the same animal model but were unable to reproduce many of the results reported by Menuet and colleagues. However, they did find a significant reduction in the number of USVs per minute at 8-9 months old compared to 5-6 months old in both Tau.P301L and wild-type mice (Trevizan-Bau et al., 2019).

These studies highlight heterogeneity in USV recordings, leading to different observations. This will be returned to in the Discussion.

##### Tau.P301S

Another well-known model was described by Scattoni and colleagues (Scattoni et al., 2010). In this study, hetero- and homozygous Tau.P301S mice were examined at postnatal days 1, 3, 5, and 7. Only the number of USVs was analysed, revealing an increase in hetero- and homozygous P301S mice at PND3, 5, and 7 compared to controls. In addition, homozygous animals produced more USVs on PND5 than their heterozygous counterparts, suggesting potential early changes in neural circuits related to vocalisations.

Together, these findings indicate that FTDP-17–linked tau mutations can alter USV production in mice, although the direction and magnitude of these effects vary across models, ages, and experimental conditions. While Tau.P301L studies report age-dependent reductions in several acoustic parameters, results remain inconsistent between laboratories, underscoring methodological variability. In contrast, Tau.P301S mice exhibit increased vocal output during early postnatal development, suggesting that tau pathology may differentially affect communication-related behaviours across developmental stages and disease progression.

### FTD-ALS models

Compared with models of PD, HD, and AD, the number of studies investigating USVs in FTD-ALS models is very sparse. This is surprising, as clear speech deficits are found relatively early after diagnosis (Di Stefano et al., 2016; Hsiung et al., 2012; Saracino et al., 2021), but may reflect a lack of mouse models or laboratories explicitly studying this condition.

#### Genetic models

##### C9BAC

The most common genetic cause of FTD and ALS is a hexanucleotide repeat expansion (HRE) in the chromosome 9 open reading frame 72 (C9orf72) gene, and expansion in the gene is associated with TDP-43 pathology (DeJesus-Hernandez et al., 2011; Renton et al., 2011).

Kahriman and colleagues used a previously established bacterial artificial chromosome transgenic C9orf72 (C9BAC) mouse model (Peters et al., 2015). The homozygous mouse has around 500 repeat motifs, and the hemizygous mouse expresses three to five copies of the transgene.

The authors did not identify significant differences in call presence, duration, amplitude and peak frequency between non-transgenic sham controls, sham C9BAC^tg/-,^ and sham C9BAC^tg/tg^ animals (Kahriman et al., 2023).

### Observed neural and circuit changes in models

In addition to changes in USVs, the included papers also described correlations with changes in neurons and circuits across the various animal models. These are further detailed below.

#### Parkinson’s Disease models

##### Alpha-synuclein overexpression models

Although neither study directly correlates USV findings to brain neurochemistry, the administration of rAAV2/5-human-𝛼-Syn into the substantia nigra (SN) significantly reduced tyrosine hydroxylase immunoreactive (TH+) neurons with 50% compared to the rAAV2/5-GFP group at 4 weeks post-injection (Gombash et al., 2013).

One, two, and six months after unilateral intrastriatal non-fibrillised recombinant 𝛼-syn and PFF injections, hyperphosphorylated 𝛼-Syn accumulations were observed ipsilaterally in the SNc, with significantly greater accumulations following PFF injection (Paumier et al., 2015). Lewy-body inclusions were also observed in areas that innervate the striatum (STR), most prominently the prefrontal, insular, cingulate, and motor cortices (layers IV–V) as well as the amygdala. The SNc also showed bilateral loss of dopamine neurons and ipsilateral loss of dopamine dendrites at 6 months after PFF injection.

The STR also showed substantial neuropathology (Paumier et al., 2015). At 2 and 6 months after intrastriatal PFF injections, there were bilateral reductions in striatal dopaminergic innervations. At 6 months post-PFF injections, there were ipsilateral diminished dopamine levels and deficits in homovanillic acid (HVA) bilateral and 3,4-dihydroxyphenylacetic acid (DOPAC) ipsilateral. Finally, there were also significant deficits in striatal dopamine turnover (HVA/DA) ipsilateral to the non-fibrilised recombinant 𝛼-syn injection site and contralateral to the PFF injection site.

Immunohistochemical findings also revealed alpha-synuclein aggregates in the Thy1-𝛼Syn mice in the brainstem and specifically the dorsal and ventral periaqueductal gray (PAG) at 5 months (Grant et al., 2014).

#### Genetic models

##### DJ1-/- model

Immunohistochemical analysis showed a significant reduction in TH+ cell bodies in the locus coeruleus (LC) of the DJ1-/- rats at 8 months of age compared to control animals (Yang et al., 2018).

##### Pink1-/- model

A wide range of neurochemical, histological, molecular, and imaging approaches were used to characterise neurological alterations in Pink1 rat models. These investigations collectively highlight region-specific changes, as well as, on occasion, contradictory changes across monoaminergic, dopaminergic, and broader neural systems.

Although classically thought of as a dopamine deficit disorder, several studies also explored serotonergic alterations. The following studies similarly defined the relative optical density of serotonin (5-HT) immunoreactivity (Broadfoot et al., 2024; Hoffmeister et al., 2024; Johnson et al., 2020). Johnson and colleagues examined 5-HT immunoreactivity in the dorsal raphe nucleus and reported no significant genotype-dependent differences at 10 months of age (Johnson et al., 2020). In contrast, Broadfoot and colleagues found reduced 5-HT immunoreactivity in the hypoglossal nucleus (12N) of 4-

month-old Pink1-/- rats (Broadfoot et al., 2024). Interestingly, Hoffmeister and colleagues reported an opposite change at 6 months of age, with wild-type rats exhibiting lower 5-HT levels in the 12N than Pink1-/- animals (Hoffmeister et al., 2024). Together, these findings suggest that serotonergic changes may be both region- and age-dependent.

Alterations in catecholaminergic systems were also examined. Using immunohistochemistry, Grant and colleagues demonstrated a reduction in TH+ neurons in the LC of 8-month-old Pink1-/- and Pink1+/-rats relative to wild-type controls (Grant et al., 2015). Notably, this reduction was significantly and positively correlated with decreased vocalisation intensity in Pink1-/- animals, indicating a potential functional link. In contrast, no genotype-dependent differences were detected in TH+ optical density in the STR or in dopaminergic cell counts within the SNc.

The same study further assessed pathological protein aggregation, revealing a pronounced accumulation of insoluble 𝛼-synuclein-positive aggregates in Pink1-/- rats across multiple regions, including the PAG, SNc, LC, STR, and nucleus ambiguus (nAmb) (Grant et al., 2015). Aggregate density was highest in the PAG, SNc, and LC. Wild-type rats showed no detectable aggregates in any of these regions, whereas Pink1+/– rats displayed a limited number of inclusions, suggesting a gene-dose effect.

Beyond monoaminergic systems, Stevenson and colleagues examined transcript and protein expression of dopaminergic, GABAergic, and opioid markers in the ventral tegmental area of 8-month-old rats (Stevenson et al., 2019). While TH and GABAergic markers were unaffected, wild-type animals exhibited significantly higher mu-opioid receptor densities than Pink1-/- rats, pointing toward altered opioid signalling within the mesolimbic circuitry.

Neurochemical profiling further revealed increased concentrations of 3-methoxy-4-hydroxyphenylglycol (MHPG), a metabolite of norepinephrine, in the STR of 8-month-old Pink1-/- rats compared with wild-type controls (Marquis et al., 2020), indicating elevated noradrenergic turnover.

At the systems level, Converse and colleagues conducted a [¹⁸F]-FDG PET imaging study to assess brain glucose metabolism in animals aged 9-12 months (Converse et al., 2024). The authors identified genotype-dependent metabolic changes in six regions, with reduced whole-brain-normalised FDG uptake in the prelimbic cortex, STR, caudate putamen, and globus pallidus, alongside increased uptake in the nAmb and posterior parietal association cortex.

Finally, transcriptomic analyses of the brainstem revealed significant molecular alterations (Lechner et al., 2022). Lechner and colleagues identified *Tuba1c* as the most upregulated gene in Pink1-/- rats of both sexes, with corresponding increases in protein expression, particularly in females. The relative mRNA quantity (as measured by real-time qPCR) of *Tuba1c* was also significantly increased in males. Furthermore, *Tuba1c* transcript levels were negatively correlated with USV top-10 peak frequency, suggesting a potential molecular link to vocal communication deficits.

In mice, Kelm-Nelson and colleagues studied neural tissue from 6-month-old animals and found no difference in the number of TH+ cells in the SN between Pink1-/- mice and controls. Also, the optical density of TH+ neurons in the STR did not differ significantly between genotypes (Kelm-Nelson et al., 2018).

### Huntington’s Disease models

#### Genetic models

##### tgHD

In this study gene expression in the STR of homozygous tgHD P10 rat pups was analysed and aberrant regulation of eight genes was found which included: angiotensinogen (*Agt*), ATPase, Ca^2+^ transport (*Atp2A2*), Forkhead box G1B (*FoxG1B*), hypocretin (orexin) receptor (*Hcrtr2*), potassium channel, member 1 (*Kcnc1*), solute carrier family 6, 3 (dopamine transporter) (*Slc6A3*), and tyrosine hydroxylase (*Th*) (Siebzehnrubl et al., 2018). In contrast to observed upregulation of *Slc6A3* and *Th*, DARPP-32, the dopamine receptor D1A, and protein kinase A (PKA) levels were significantly down-regulated in the STR. Western blot analyses of striatal protein extracts also confirmed a reduction in DARPP-32 protein levels.

An additional autoradiography experiment showed decreased density of the dopaminergic D1A receptor and the NMDA receptor (NMDAR) in tgHD as compared to wild-type rats (Siebzehnrubl et al., 2018). Western blot analyses showed down-regulation of NR1, NR2B and NR2C NMDAR subunits. Immunohistochemical analysis of P10 tgHD striata confirmed reduced expression of DARPP32 and TH. This paper also characterised the tgHD rat from a neurodevelopmental perspective and found a significant reduction in the content of Dcx+ neurons, expressed by neuronal precursors, in the rostral migratory stream (RMS) and olfactory bulb (OB) (Siebzehnrubl et al., 2018), corroborating similar findings of a clear developmental role for htt (Barnat et al., 2020; Barnat et al., 2017; Braz et al., 2022).

##### BACHD

Postmortem studies of hypothalamic tissue from individuals with HD have revealed a marked loss of the neuropeptides oxytocin and vasopressin, which are key regulators of emotional and social behaviour (Gabery et al., 2015; Gabery et al., 2010). Because the dynamic balance between these neuropeptides is crucial for normal socio-emotional functioning, disruptions in their interplay may contribute significantly to the neuropsychiatric manifestations of the disease. Therefore, Cheong and colleagues focused on oxytocin and vasopressin to better understand their role in HD (Cheong et al., 2020).

The concentrations of oxytocin and vasopressin were analysed in the hypothalamus, amygdala, pituitary, and plasma of 2-, 6-, 12-, and 15-month-old mice. Firstly, oxytocin was increased in the hypothalamus of 12-month-old mice and reduced in the pituitary at 6 months of age in female BACHD mice and plasma at 6 and 15 months of age in BACHD compared to age-matched wild-type animals. Secondly, plasma vasopressin levels were increased in BACHD mice compared to wild-type mice at 2, 6, and 12 months of age (Cheong et al., 2020). This data shows an imbalance of the oxytocin-vasopressin system.

### Alzheimer’s Disease models

#### Neurotoxin models

Sap-lesioned rats had reduced choline acetyltransferase (ChAT) immunoreactivity in the hippocampus and cortex, whereas caspase-3 activity was increased in the hippocampus and cortex compared to controls, suggesting changes in the regulation of apoptosis (Cutuli et al., 2013).

#### Genetic models

##### McGill-R-Thy1-APP

Petrasek and colleagues performed RT qPCR on rat brains at 10 months of age, sacrificed at Zeitgeber Time (ZT) 3, three hours after the onset of light, to look at the expression of circadian clock genes *Bmal1, Nr1d1,* and *Prok2,* and found that the relative expression of *Bmal1* and *Prok2* was significantly increased in the parietal cortex and cerebellum, and parietal cortex and hippocampus, respectively, of McGill-R-Thy1-APP compared to wild-type rats (Petrasek et al., 2018).

### Models with tauopathy

#### Genetic models

##### Tau.P301L

Neurons positive for tau protein phosphorylated at residues threonine 212 and serine 214 (AT100+) were strongly expressed in the old age group (8-10 months old) in the PAG and intermediate in other brainstem structures, such as the nucleus reticularis gigantocellularis (NRA) and kölliker-fuse (KF) nucleus (Menuet et al., 2011). Neurons positive for tau protein phosphorylated at serine 202 and threonine 205 (AT8+) were also observed in the older age group, with high expression in the brainstem, including the PAG, KF and NRA.

Trevizan-Bau and colleagues investigated the relationship between tauopathy in the PAG, KF and NRA and the observed USV phenotype. The phenotypes included hypoactive (0-35 USVs per min), average (90-205 USVs per min), or hyperactive (>260 USVs per min) USV production. However, no relationship was found. (Trevizan-Bau et al., 2019).

##### Tau.P301S

Human tau expression was analysed from postnatal day 1 until 14. Transgenic human tau was detected from PND1 in both homozygous and heterozygous P301S tau transgenic mice. Expression of human transgenic tau steadily increased with age in both the brain and the spinal cord, and was higher in homozygotes than in heterozygotes (Scattoni et al., 2010).

### FTD-ALS models

#### Genetic models

##### C9BAC

Although Kahriman and colleagues found no differences in USV parameters between C9BAC^tg/tg^, C9BAC^tg/-,^ and controls, these animals showed significant neural effects (Kahriman et al., 2023). First, at 15 months of age, sham-operated C9BAC^tg/-^ and C9BAC^tg/tg^ mice exhibited a significant loss of SMI-312 cortical staining and SMO-312-stained axon profiles. Moreover, this effect was even more pronounced in C9BAC^tg/tg^ than in hemizygous animals. Additionally, the postsynaptic marker PSD-95 and presynaptic marker synaptophysin were both significantly reduced in sham-operated hemi- and homozygous mice compared to controls.

## Discussion

Here, we have provided an overview of the current literature on USV generation in rodent models of neurodegenerative disease. The most commonly used models for studying the effects of neurodegeneration on USV production are the Pink1-/- rat model and the R6/1 mouse model; therefore, PD and HD are the most examined diseases. For a few models, the review showed consistent findings for USV features across studies. In Pink1-/- rats, the average bandwidth and peak frequency of 50-kHz FM calls were significantly reduced. Haloperidol-treated rats and R6/1 transgenic mice exhibited a significant reduction in the number of calls. In BACHD mice, both total calling time and call number were consistently reduced across studies. Moreover, the number of calls was reduced across species and neurodegenerative models, including PD, HD, AD and tauopathies.

The study of various neurodegenerative models also provided insight into the roles of distinct brain structures and neural circuits in the production of USVs. In PD, studies using the 6-OHDA neurotoxic drug show that dopamine in the nigrostriatal pathway, specifically in structures such as the SNc, plays an important role in USV production. Moreover, the vocal repertoire was more severely affected after antagonising the function of D1 and D2 receptors. Additionally, vocal impairment also occurs when TH is reduced in the LC, as shown in the DJ1-/- and Pink-/- model. Besides dopamine, other systems were also impaired. For example, the VTA exhibited reduced mu-opioid receptor levels in Pink1-/- rats. The striatum also showed elevated noradrenergic turnover in Pink-/- mice and reductions in striatal dopaminergic innervations in alpha-synuclein overexpressing models.

However, the role of tyrosine hydroxylase (TH) needs further investigation. For example, TH in the LC is not affected at 4 months (Broadfoot et al., 2024), but is at 8 months in the same Pink-/- rat model (Grant et al., 2015). However, Broadfoot and colleagues already reported a reduction of peak frequency at 2 months of age.

An overall observation among studies is that neuropathology is investigated after the first changes in vocalisations occur. Therefore, the earliest neural circuits underlying vocal deficits in neurodegeneration remain underexplored.

Some studies also examined serotonergic alterations in PD models. This idea arose from the observation that vocal and swallowing deficits show little improvement with dopamine replacement (Brabenec et al., 2017; Menezes & Melo, 2009), unlike limb motor symptoms, suggesting that their underlying mechanisms are at least partly extra-dopaminergic (Rudisch, Krasko, Burdick, et al., 2023). Evidence points to early disruption of brainstem serotonergic and noradrenergic systems that innervate cranial sensory-motor nuclei, with several studies reporting 5-HT and noradrenergic degeneration preceding classic motor onset (Hoffmeister et al., 2021; Lewitt, 2012; Wilson et al., 2019; Wohr et al., 2015). Notably, tongue-strengthening exercise can reverse age-related loss of 5-HT input to the 12N (Behan et al., 2012), and serotonergic modulation strongly affects vocalisation (Wohr et al., 2015), highlighting 5-HT dysfunction in the hypoglossal system as a promising target for understanding and improving voice and swallowing impairments in PD. However, serotonergic levels in the 12N change across age in Pink1-/- rats, while USV features, such as peak frequency, showed a constant decline over time(Broadfoot et al., 2024; Hoffmeister et al., 2024). This shows that further research is needed to identify the role of extra-catecholaminergic neurotransmitter systems affected in neurodegeneration.

In HD, studies in tgHD pups show dysregulation of dopaminergic and glutamatergic signalling at the stage when USV production declines. This parallels imaging findings in asymptomatic HD gene carriers (Ginovart et al., 1997; Weeks et al., 1996), which reveal reductions in striatal D1 and D2 receptors, a pattern further supported by molecular analyses of postmortem HD brains (Glass et al., 2000). In contrast, studies found a loss of NMDA receptors in the striatum of pre-symptomatic and symptomatic cases (Fan & Raymond, 2007). This may reflect disrupted interactions between glutamate and dopamine receptors and their roles in USVs.

This review shows that dopamine receptor antagonists, such as haloperidol, markedly affect USV production. It is important to note that haloperidol is also used to treat agitation and aggression in people with dementia (Fox et al., 2025). This effect is also evident in rats treated with haloperidol, which showed a reduction in the duration and number of aversive 22-kHz calls during foot shock presentation (Colombo et al., 2013). These results underline the need to consider both psychopharmacological treatments and neuropathological factors when looking at speech changes in people with dementia.

In taupathology models, hyperphosphorylated tau was present in key brainstem and spinal cord structures involved in USV production. All taumodels showed affected USV production, and, interestingly, vocal dysfunction in the Tau.P301L mice preceded taupathology and was not linked to the number of USVs produced. This indicates that earlier neural alterations underlie these early vocal changes.

Within the literature, specific disease models yield contrasting findings. The contrasting findings regarding intensity in Pink1-/- rats might be attributed to differences in data reporting and to sex differences. Firstly, Grant and colleagues, and Marquis and colleagues, reported main effects of genotype, collapsed across testing age, whereas Lechner and colleagues, and Johnson and colleagues, examined age-specific differences (Grant et al., 2015; Johnson et al., 2020; Lechner et al., 2022; Marquis et al., 2020). Secondly, Lechner and colleagues studied both sexes and genotypes in the same testing cohort and showed that female rats produced FM calls with significantly reduced intensity than male rats, regardless of genotype (Lechner et al., 2022). Thus, it is important to consider sex and age differences when reporting USV data.

Also, for the Tau.P301L mice, different results were reported. Different experimental conditions can partially explain these differences. Menuet and colleagues recorded USVs during unfamiliar social interaction between one male and two female mice (Menuet et al., 2011). In contrast, Trevizan-Bau and colleagues recorded USVs from one male and one female mouse during courtship (Trevizan-Bau et al., 2019). Moreover, vocalisation is a complex behaviour influenced by numerous innate and environmental factors (Chabout et al., 2012; Kikusui et al., 2011). For example, subtle differences in housing conditions can affect USV production in genetically homogeneous mice (Brenes et al., 2016). Additionally, the female oestrus phase strongly influences a male rodent’s motivation to emit USVs (Barthelemy et al., 2004; Hanson & Hurley, 2012; McGinnis & Vakulenko, 2003). Therefore, assessing the oestrous cycle phase is crucial for replicating results. Nevertheless, the USV pattern and the amount of USV activity reported by Trevizan-Bau and colleagues were consistent with those reported by Menuet and colleagues.

### Quantitative comparison between rodent models and patients

Despite the vast differences between species, some similarities could be drawn between USV changes observed in rodent models and motor speech abnormalities in patients with neurodegenerative disease. Critically, the possibility to explore such comparisons is heavily rooted in the shift from qualitative descriptions to quantitative, reproducible metrics implemented for semi-automated digital speech analysis.

Across rodent models of PD, HD, and tauopathy, reduced call numbers and call durations are consistently observed. Although lacking disease specificity, a parallel between this USV metric may be drawn with the observations that speech rate decreases or pause durations increase across patients with PD, HD and PSP (Berthier et al., 2026; Coppieters et al., 2024; Nunes et al., 2024) In rodent models of PD and HD, a decrease in the peak USV frequency was also robustly demonstrated, whereas more data is needed in tauopathy models. In a similar vein, in patients with PD and HD, aprosody is a common finding (Diehl et al., 2019; Xu et al., 2025)

Within the spectrum of PD and related disorders such as HD and PSP, direct comparisons between digital speech markers of diseases are limited. However, some studies described shared abnormalities across these movement disorders, e.g., decreases in voiced time compared to controls, combined with more pronounced decreases in articulatory rate in PSP compared to PD (Kang et al., 2025; Kang et al., 2023). Similar studies, including head-to-head comparisons in rodent models, that document the relative magnitude of USV changes are currently lacking.

Both Petrasek and colleagues and Pietropaolo and colleagues studied amyloid-beta models in rats and mice, respectively, and linked vocal alterations to social deficits. Both animal models showed distinct behavioural profiles during social interaction, accompanied by distinct USV profiles compared with controls (Petrasek et al., 2018; Pietropaolo et al., 2012). The social alterations observed in AD models parallel those reported in patients, including behavioural changes such as apathy, withdrawal or sometimes disinhibition, as well as changes in social cognition (Jicha & Carr, 2010; Mohs et al., 2000; Rankin et al., 2008). Additional alterations in behaviour were shown by Cutuli and colleagues, who showed that Sap lesions in rats reduced the production of fear-associated USVs in response to foot shock, while the freezing response remained unchanged (Cutuli et al., 2013). This dissociation between freezing response and USV production in rats with cholinergic lesions is consistent with other studies (Frick et al., 2004), and they hypothesise that this may be linked to coping behaviour.

## Limitations

Certain limitations should be taken into consideration when interpreting the findings of this review. First, not all studies that paired a male with a female animal systematically controlled for or reported the female’s oestrus phase, which can influence the observed vocal repertoire. In addition, there exists considerable heterogeneity in the paradigms used to elicit USVs, including mating and social interaction paradigms, which are associated with distinct affective states and may therefore differentially impact vocal output. This methodological variability is further exacerbated by the wide diversity in the acoustic parameters extracted and analysed across studies, which limits direct comparisons. Secondly, there is a general lack of recordings from female rodents, alongside a broad age range in which USVs are recorded, both of which limit the ability to draw firm conclusions regarding sex- and age-related effects. Thirdly, the literature is mainly focused on disorders with dominant motor symptoms, such as PD and HD, rather than those characterised by broader, more cognitive symptom profiles, such as AD and FTD, potentially biasing the overall conclusions. Fourthly, the use of different rodent strains across studies introduces additional variability that may contribute to inconsistent results. Finally, in most animal models, only a limited number of studies were available, making it difficult to identify clear, consistent patterns. Collectively, these limitations highlight the need for more standardised methodologies.

## Conclusion

In summary, this systematic review demonstrates that USVs constitute a sensitive and biologically relevant readout of neurodegenerative processes in rodent models. Across PD, HD, AD and tauopathy models, convergent alterations in USV production emerge, most notably reductions in call number, call duration, bandwidth and frequency, often appearing early in disease progression and, in some cases, preceding overt neuropathology or classical motor deficits. These vocal changes reflect dysfunction across distributed neural circuits spanning the brainstem, midbrain, and forebrain, implicating not only dopaminergic systems but also other neuromodulatory and peptidergic pathways. At the same time, substantial heterogeneity in elicitation paradigms, acoustic metrics, ages, sexes, and species limits cross-study comparability and disease specificity. Importantly, neural correlates are frequently examined after behavioural deficits have emerged, leaving the earliest circuit and molecular events underlying vocal dysfunction largely unexplored. Future studies will benefit from greater methodological standardisation, longitudinal designs spanning prodromal stages, systematic inclusion of females, and integrated circuit-level analyses. Addressing these gaps will be critical to establishing USVs as quantitative, cross-species biomarkers that bridge preclinical models and human speech and communication deficits, thereby advancing early detection and mechanistic understanding of neurodegenerative disease.

## Supplementary materials

### S1

| PubMed |
| --- |
| ("Murinae"[MeSH] OR rat OR rats OR mouse OR mice) AND ("Dementia"[MeSH] OR "Neurodegenerative Diseases"[MeSH] OR dement* OR alzheimer* OR huntingt* OR neurodegenerati* OR "degenerative disease*" OR tauopathy* OR tau OR parkinson* OR frontotemporal* OR lewy* OR pick* OR "primary progressive aphasia" OR FTLD OR FTD OR ALSFTD) AND ("Ultrasonics"[MeSH] OR "Vocalization, Animal"[MeSH] OR "Ultrasonic Waves"[MeSH] OR "Animal Communication" [MeSH] OR ultrasonic* OR ultrasound* OR vocalization* OR vocalisation* OR USV OR calls OR "animal communication*" OR "animal sound")<br>((#2) NOT ("review"[Publication Type])) NOT ("systematic review"[Publication Type]) |
| Web of Science |
| ALL=((rat OR rats OR mouse OR mice) AND (dement* OR alzheimer* OR huntingt* OR neurodegenerati* OR "degenerative disease*" OR tauopathy* OR tau OR parkinson* OR frontotemporal* OR lewy* OR pick* OR "primary progressive aphasia" OR FTLD OR FTD OR ALSFTD) AND (ultrasonic* OR ultrasound* OR vocalization* OR vocalisation* OR USV OR calls OR "animal communication*"))<br>(#5) NOT DT=(Review) |
| Embase |
| ('murine'/exp OR rat OR rats OR mouse OR mice) AND ('dementia'/exp OR 'degenerative disease'/exp OR dement* OR alzheimer* OR huntingt* OR neurodegenerati* OR "degenerative disease*" OR tauopathy* OR tau OR parkinson* OR frontotemporal* OR lewy* OR pick* OR "primary progressive aphasia" OR FTLD OR FTD OR ALSFTD) AND ('ultrasound'/exp OR 'vocalization'/exp OR 'animal communication'/exp OR ultrasonic* OR ultrasound* OR vocalization* OR vocalisation* OR USV OR calls OR "animal communication*" OR "animal sound*") NOT review:it |

## Acknowledgements

We thank Barbara Lejeune for helping us to set up the appropriate search strategy.

